# Lipid-rich ascites reprograms T cell lipid metabolic transcriptome to drive dysfunction

**DOI:** 10.64898/2026.03.11.711101

**Authors:** Peter Kok-Ting Wan, Gulsah Albayrak, Luke Furtado O’Mahony, Kerry Fisher, Leonard W Seymour

## Abstract

**Purpose:** Bispecific T cell engagers (BiTEs) have recently been approved as a locoregional immunotherapy for malignant ascites. Although ascites is recognised as a lipid-rich, immunosuppressive environment, the mechanisms by which ascites, particularly its lipid components, suppress antitumour immunity remain poorly understood. Here, we investigated the impact of ascites-associated lipids on T cell immunosuppression and assessed whether lipid modulation could enhance the efficacy of BiTE therapy.

**Experimental Design:** Transcriptomic profiling was performed on T cells treated with acellular ascites fluid to identify gene expression signatures associated with ascites exposure. Functional assays were conducted to evaluate the effects of ascites-associated lipids on T cell activation and cytotoxicity. In parallel, T cells were cocultured with ovarian cancer cells and EpCAM-targeting BiTEs in the presence or absence of a lipid-removal agent to assess how lipid depletion affected BiTE efficacy.

**Results:** T cells exposed to acellular ascites fluid exhibited an enriched transcriptomic signature associated with cholesterol efflux and incomplete fatty acid oxidation, which are metabolic features often found in exhausted T cells. These alterations converged on a metabolically imbalanced state linked to impaired plasma membrane signalling. Lipid removal from ascites selectively rescued CD137 expression but not CD25, and restored BiTE-mediated cytotoxicity, suggesting a differential impact of lipid metabolism on TCR complex-dependent versus cytokine-driven activation pathways.

**Conclusions:** These findings identified lipid as a driver for T cell dysfunction in ovarian cancer ascites. Removal of ascites lipids restored T cell activation and augmented BiTE-mediated cytotoxicity, supporting a combination approach to potentiate BiTE therapy in malignant ascites.

**Translational Relevance:** Malignant ascites represents a lipid-rich, immunosuppressive tumour microenvironment that is increasingly targeted by emerging T cell-based therapies. Although EpCAM-targeting bispecific T cell engagers (BiTEs) have recently been approved for malignant ascites and multiple similar BiTEs are in clinical development, the mechanisms by which ascites impairs T cell function and potentially limits therapeutic efficacy remain poorly understood. Using patient-derived ascites throughout, this study demonstrated that lipid metabolic reprogramming, rather than immune checkpoint upregulation, was a driver of T cell dysfunction. Importantly, we demonstrated that lipid removal from ascites rescued T cell function and restored BiTE efficacy, identifying a targetable metabolic barrier to immunotherapy. While EpCAM was used as a proof-of-concept target, we anticipate the metabolic insights and therapeutic strategies identified here will be equally applicable to other BiTE and CAR-T platforms, supporting a new combination approach for the treatment of malignant ascites.

## INTRODUCTION

Ovarian cancer remains the most lethal gynaecological malignancy, with most patients diagnosed at an advanced stage and five-year survival rates remaining below 30% despite advances in surgery and chemotherapy (1). Advanced ovarian cancer readily metastasises to the omentum, peritoneum, and other abdominal organs (2). As metastasis progresses, many patients develop large volumes of ascites that foster immunosuppression and tumour progression (3,4). Although the immunosuppressive effects of ascites on T cells are well recognised, where they demonstrated impaired proliferation, reduced cytokine secretion, and attenuated cytotoxicity, the mechanisms driving this dysfunction remain poorly understood (5–7).

Recent work highlights cellular metabolism could be a central regulator of T cell fate and function (8–10). Ascites is enriched in lipids, and altered lipid metabolism has been linked to T cell dysfunction in solid tumours (11). Excessive lipid exposure has been reported to induce oxidative stress through lipid peroxidation, disrupt mitochondrial integrity, and reshape transcriptional cascades that govern T cell proliferation and effector function (12,13). Altered fatty acid and cholesterol metabolism could further impair membrane organisation, signalling competence, and metabolic flexibility, ultimately limiting T cell persistence and skewing their fate toward dysfunctional phenotypes within tumour microenvironments (13–16). It is increasingly recognised that a better understanding of lipid-driven T cell dysfunction in ascites could provide essential insights into tumour immune evasion and guide the development of more effective therapeutic strategies.

In this study, we integrated transcriptomic and functional analyses to characterise the lipid metabolic states of different T cell subsets and dissect how ascites altered T cell metabolism. We found that exhausted T cells exhibited transcriptomic signatures consistent with reduced cholesterol retention and impaired fatty acid oxidation, suggesting a metabolically imbalanced state, whereas regulatory T cells displayed a lipid-adapted metabolism. Importantly, we showed that exposure to acellular ascites reprogrammed otherwise functional T cells into a metabolically imbalanced, signalling-impaired state, in which they adopted a lipid transcriptomic profile resembling that of exhausted T cells but without broad immune checkpoint upregulation. We also demonstrated that lipid depletion restored T cell activation and augmented bispecific T cell engager (BiTE) activity, implicating lipid metabolism as a key factor in T cell dysfunction in ovarian cancer ascites.

## MATERIAL AND METHODS

### Cell culture and primary sample processing

For *in vitro* cell culture, ovarian cancer HeyA8 cells were maintained in Roswell Park Memorial Institute medium (RPMI; Sigma-Aldrich) supplemented with 10% heat-inactivated fetal bovine serum (FBS). The cell line was obtained from American Type Cell Collection (ATCC). Cells were cultured in a humidified incubator at 37 °C with 5% CO₂ unless otherwise specified. All cell lines tested negative for mycoplasma using the MycoAlert Mycoplasma Detection Kit (Lonza).

Primary human malignant ascites samples were obtained from Churchill Hospital (Oxford, UK) and immediately separated into cellular and fluid fractions by two rounds of centrifugation at 400 × g. The cellular fractions were subsequently treated with red blood cell lysis buffer (Qiagen). Details of the clinical samples used in this study are provided (fig S1).

PBMCs were isolated from human leukocyte cones obtained from consented healthy volunteers (NHS Blood and Transfusion Service, Oxford, UK) by density-gradient centrifugation with Ficoll-Paque Plus (GE Healthcare). CD3⁺ T cells were subsequently purified from PBMCs using the Pan T Cell Isolation Kit (Miltenyi Biotech).

For *in vitro* ascites cultures, PBMC-derived T cells from healthy donors were cultured with acellular ascites fluid. Where required, lipids were depleted from ascites using the adsorption and clarification reagent, Cleanascites (Biotech Support Group).

### Coculture experiment

CD3^+^ T cells were isolated from human PBMCs, and cocultured with HeyA8 cells at a 5:1 ratio in 2% FBS-supplemented RPMI, acellular ascites fluid mixed (50% v/v) with FBS-supplemented RPMI, or lipid-depleted acellular ascites fluid mixed (50% v/v) with FBS-supplemented RPMI, in the presence of either EpCAM BiTE or Control BiTE. Where indicated, CD3/CD28 Dynabeads (Thermo Fisher) were used to induce T cell activation in the absence of HeyA8 cells. After 72 hours, T cells were harvested, stained with fluorescently labelled antibodies, and analysed by flow cytometry. Target cell cytotoxicity was assessed using the XTT Cell Proliferation Assay II (Roche).

### RNA sequencing and computational analysis

Purified T cells were resuspended at a density of 1 × 10⁶ cells/mL in complete RPMI medium without additional stimulation and rested for 24 hours. Cells were then harvested, and total RNA was extracted using TRIzol (Invitrogen) as described previously (17). RNA was eluted in RNase-free water, and quality was confirmed by NanoDrop spectrophotometry (A260/A280 > 1.8). Libraries were prepared using a standard poly(A) selection workflow. Sequencing was carried out on the Illumina NovaSeq X Plus with at least 20 million reads per sample (Genewiz).

RNA-seq data were processed starting from aligned BAM files. Gene-level counts were quantified using *featureCounts* with the GRCh38.110 GTF annotation. Ensembl gene identifiers were converted to HGNC gene symbols using *biomaRt*. Low-quality features were removed prior to downstream analysis. The filtered count matrix was used to construct a DESeq2 dataset, with treatment condition specified as the design factor. Differential expression analysis was performed, and significantly differentially expressed genes (DEGs) were defined as those with adjusted P value (FDR) < 0.05 and absolute log₂ fold change > 1. Variance-stabilising transformation (VST) was applied to normalise expression values for visualisation. Heatmaps of the top 50 DEGs were generated using *pheatmap*, and volcano plots were produced using *ggplot2* and *EnhancedVolcano*. Gene Set Enrichment Analysis (GSEA) was performed across MSigDB collections (Hallmark and C5) using ranked gene lists based on log₂ fold change values, with enrichment visualised by dot plots and enrichment curves. All analyses were performed in R (version 4.4.1).

### Analysis of single-cell RNA sequencing data

Processed scRNA-seq data from the Zheng et al. dataset were obtained from Mendeley Data (18). T cells were extracted from the processed data, normalised, scaled, and visualised using uniform manifold approximation and projection (UMAP). Cell clusters were identified with the Seurat functions *FindNeighbors* and *FindClusters* and annotated based on canonical cell markers. Curated gene signatures were scored using *AddModuleScore* and compared across subclusters.

### Flow cytometry

Cells were plated in V-bottom 96-well plates (Corning) in MACS staining buffer and washed twice with the same buffer before incubation with human FcR blocking reagent (Miltenyi Biotech) for 15 min at room temperature. For surface staining of CD3 (RRID:AB_314052), CD4 (RRID:AB_571951), CD8 (RRID:AB_314116), CD11b (RRID:AB_2562020), CD25 (RRID:AB_11218989), CD56 (RRID:AB_2566060), CD137 (RRID:AB_2563830), EpCAM (RRID:AB_756082), and PD-1 (RRID:AB_2159324), cells were incubated with the corresponding antibodies at a 1:200 dilution in MACS buffer. Stained cells were then fixed with 10% formalin, and target expression was analysed on an Attune NxT Flow Cytometer (Thermo Fisher).

For intracellular neutral lipid staining, cells were fixed and permeabilised as described, then resuspended in 100 µl of LipidTOX Green neutral lipid stain (Invitrogen) diluted 1:2000 in PBS (19). Samples were incubated for 30 min on ice, followed by resuspension in 200 µL MACS buffer before being analysed on an Attune NxT Flow Cytometer.

### Ethics for clinical samples

Primary ovarian cancer ascites were obtained from patients at Churchill Hospital (Oxford, UK) with informed consent. Ethical approval was granted by the Research Ethics Committee of the Oxford Centre for Histopathology Research (Reference ORB 20/A013).

### Statistics

Flow cytometry data were analysed using FlowJo, and all other data analyses were performed with GraphPad Prism (GraphPad Software). A t-test was used for pairwise comparisons, and two-way ANOVA with Šídák’s post hoc test was applied for grouped datasets with multiple variables. All experiments were performed in triplicate and results are presented as mean ± s.d., unless otherwise stated. Significance is indicated as follows: *P < 0.05; **P < 0.01; ***P < 0.001; ****P < 0.0001; NS, not significant.

## RESULTS

### Ascites T cells do not express high levels of immune checkpoints

To characterise the immune landscape in ovarian ascites, we analysed 13 ascites harvested from ovarian cancer patients. T cells (mean = 41.6%) were the dominant cell type in ovarian cancer ascites, with CD4⁺ T cells (mean = 27.2%) representing the major subtype (Fig 1A). CD8⁺ T cells and NK cells each contributed approximately 13-14%, such that lymphoid cells collectively accounted for more than half of the ascites population. CD11b^+^ myeloid cells (mean = 19.3%) and cancer cells (mean = 26%), as denoted by EpCAM expression, were also detected, suggesting a heterogeneous cell population in the ascites niche.

**Fig 1.**
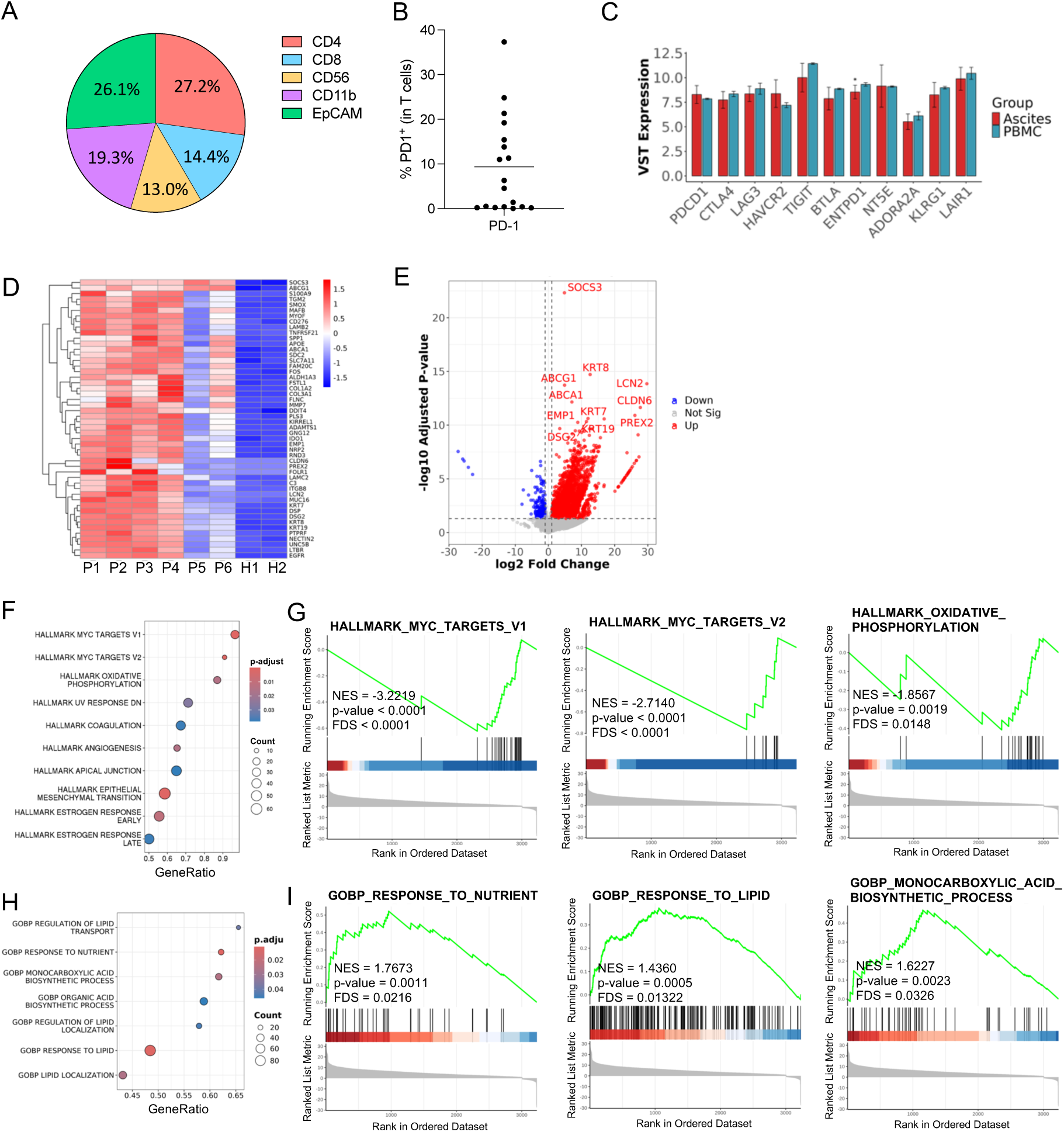
T cells in ovarian cancer ascites show limited immune checkpoint expression but are transcriptionally programmed for lipid and nutrient adaptation. (A and B) Ascites were collected from patients with ovarian cancer and analysed by flow cytometry. Relative proportions of lymphocytes, myeloid cells, and cancer cells, *n* = 13 (A). PD-1 expression on T cells, *n* = 18 (B). (C to I) Transcriptomic profiling of T cells isolated from ascites and PBMCs was performed by bulk RNA sequencing (ascites, *n* = 6; PBMCs, *n* = 2). Normalised expression of inhibitory checkpoint molecules in T cells from ascites and PBMCs (C). Heatmap (D) and volcano plot (E) showing the top differentially expressed genes between ascites and PBMCs. Dot plot of enriched pathways identified by GSEA using MSigDB Hallmark gene sets, with the top ten pathways ranked by gene ratio (F). Enrichment plots for Myc targets and oxidative phosphorylation (G). Dot plot of enriched metabolism-related pathways identified by GSEA using MSigDB C5 (Gene Ontology) gene sets (H). Enrichment plots for nutrient response, lipid response, and monocarboxylic acid biosynthesis (I). The normalised enrichment score (NES), p-values, and false discovery rate (FDR) were shown for each pathway.

Little is understood about the mechanisms by which ascites induces T cell dysfunction, even though T cells are the dominant immune cell type. We initially hypothesised that T cell dysfunction might be driven by an upregulation of inhibitory checkpoints in ascites. However, analysis of 18 ovarian ascites samples revealed that PD-1 expression on T cells was low (mean = 9.4%) (Fig 1B).

To further validate the immune profile of ascites T cells, we performed RNA-sequencing (RNA-seq) on T cells isolated from ovarian cancer ascites and compared them with healthy peripheral blood mononuclear cells (PBMCs). Ascites T cells showed no significant changes in the expression of most key genes involved in immune inhibition, including *PDCD1* (PD-1), *CTLA4*, *LAG3*, *HAVCR2* (TIM-3) and *TIGIT*, suggesting that inhibitory immune receptors may not be a primary factor driving T cell dysfunction (Fig 1C).

### Ascites T cells are programmed to respond to lipid and nutrient

Analysis of the top differentially expressed genes in ascites-derived T cells revealed enrichment of metabolic adaptation and stress-response pathways (Fig 1, D and E). Upregulation of the lipid transporters *ABCG1* and *ABCA1*, together with the lipid carrier *APOE*, indicated enhanced cholesterol efflux and altered lipid handling. The catabolic enzyme *SMOX*, which degrades spermine, suggested activation of polyamine turnover linked to oxidative stress (20). In addition, elevated expression of *DDIT4*, an mTOR inhibitor, and *SLC7A11*, a cystine-glutamate exchanger supporting glutathione synthesis, pointed to adaptations to metabolic stress (21). In parallel, the cytokine signalling inhibitor *SOCS3* was strongly induced, suggesting the T cells were immunosuppressed.

We next examined whether ascites-derived T cells displayed altered gene signatures. Gene set enrichment analysis (GSEA) of “Hallmark” pathways revealed that Myc signalling and oxidative phosphorylation were significantly downregulated in ascites T cells (Fig 1, F and G), suggesting impaired Myc-driven transcription and reduced mitochondrial function at the transcriptomic level, consistent with diminished proliferative potential. In contrast, ontologies associated with nutrient and lipid responses, as well as monocarboxylic acid biosynthesis, were positively enriched, indicating a shift toward metabolic adaptation (Fig 1, H and I). These signatures likely reflected the composition of the ascites microenvironment, in which T cells adjusted to atypical nutrient availability and stress, prioritising metabolic flexibility and survival over proliferation and effector activity.

### Exhausted T cells and regulatory T cells adapt to a high lipid metabolism

To characterise the metabolic profile of T cell subtypes, we analysed a publicly available single-cell RNA sequencing (scRNA-seq) dataset comprising 10 ascites and 6 PBMC samples from ovarian cancer patients (18). Unsupervised clustering identified three CD4⁺ subsets (naive, central memory, and regulatory T cells), four CD8⁺ subsets (naive, effector memory, effector, and exhausted), as well as mucosal-associated invariant T cells (MAIT) and γδ T cells (Fig 2, A and B). Comparison of ascites- and PBMC-derived populations revealed that ascites has a higher abundance of memory, effector, exhausted, and regulatory T cells (Fig 2C).

**Fig 2.**
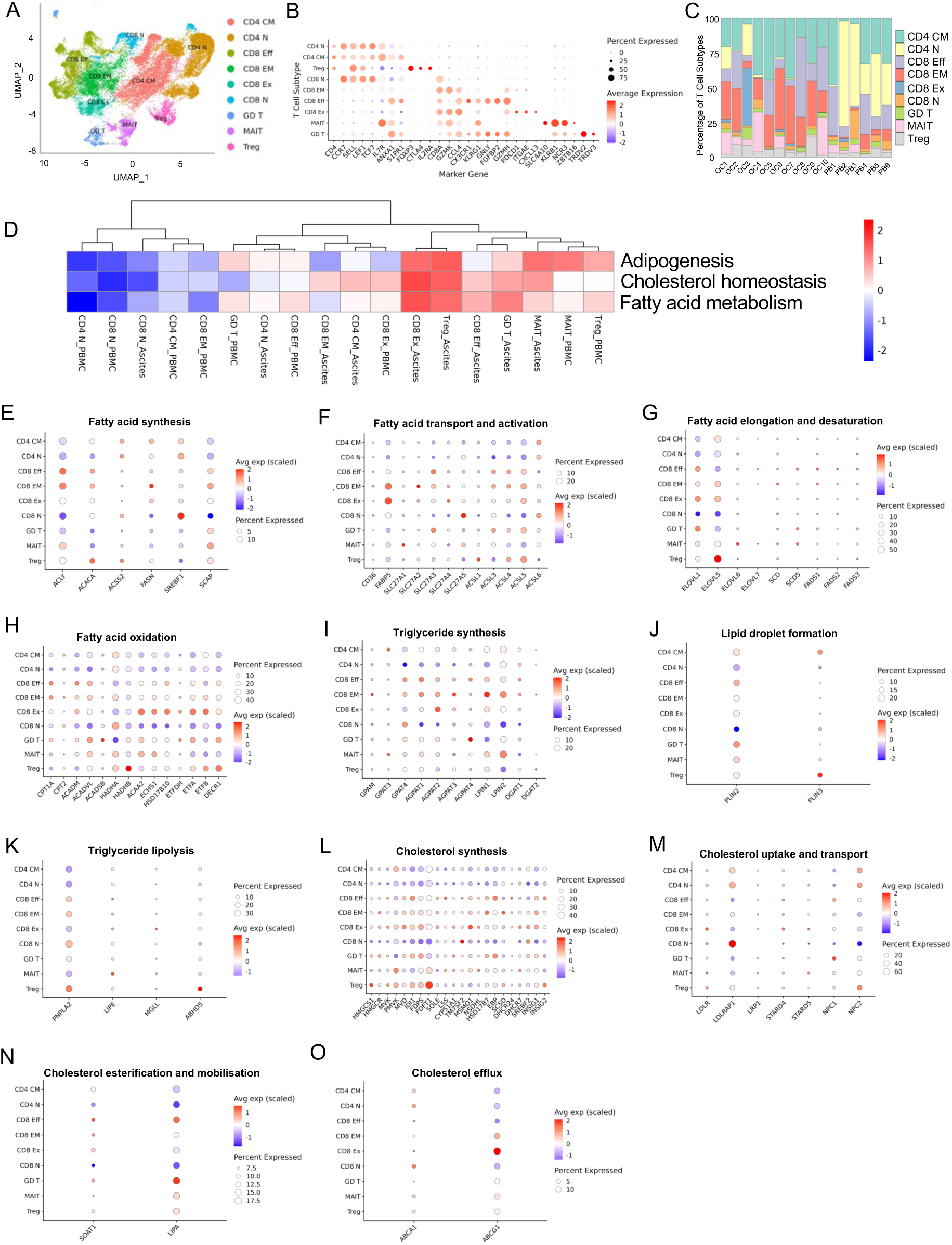
Distinct T cell subsets in ascites adopt differential lipid metabolic programmes. Transcriptomes of T cells from ovarian cancer ascites (*n* = 10) and PBMCs (*n* = 6) were analysed by single-cell RNA sequencing (scRNA-seq). (A) UMAP plot showing nine T cell subsets, with each dot representing a single cell coloured by cluster. (B) Dot plot showing the expression of selected genes across clusters. (C) Relative proportions of T cell subsets in ascites and PBMCs. (D) Heatmap of lipid metabolism-related Hallmark gene set scores across T cell subsets from ascites and PBMCs. (E to O) Dot plots showing expression of key genes involved in fatty acid synthesis (E), fatty acid transport and activation (F), fatty acid elongation and desaturation (G), fatty acid oxidation (H), triglyceride synthesis (I), lipid droplet formation (J), triglyceride lipolysis (K), cholesterol synthesis (L), cholesterol uptake and transport (M), cholesterol esterification and mobilisation (N), and cholesterol efflux (O) in ascites-derived T cells.

We next asked whether different T cell subtypes in ascites adopted similar lipid transcriptomic profiles. Surprisingly, even within the same environment, subsets exhibited distinct profiles (Fig 2D). Effector populations, particularly exhausted T cells, and regulatory T cells, showed strong signature enrichment of adipogenesis, fatty acid metabolism, and cholesterol homeostasis, whereas naive CD4⁺ and CD8⁺ naive T cells consistently displayed the lowest lipid metabolic activity.

Notably, exhausted T cells were the only subset that showed a strong cholesterol homeostasis signature in PBMCs, and this was further upregulated in ascites, suggesting that altered cholesterol regulation may represent a distinct metabolic feature predisposing T cells to exhaustion. Other effector, memory, and regulatory T cells also displayed cholesterol homeostasis enrichment in ascites, though at lower levels, implying that disrupted cholesterol metabolism may be a broader feature of ascites-driven dysfunction.

### Exhausted T cells exhibit incomplete fatty acid oxidation and accumulation of intermediates

Both exhausted T cells and regulatory T cells displayed strong lipid metabolic signatures in ascites, but these enrichment scores only captured broad pathway activity. We hypothesised that these subsets utilised lipids through distinct routes: exhausted T cells became dysfunctional due to incomplete lipid utilisation, whereas regulatory T cells exploited the lipid-rich environment to sustain their growth and immunosuppressive function. We therefore examined the key genes controlling fatty acid, triglyceride, and cholesterol metabolism.

Fatty acid metabolism was first categorised into four key phases: (i) biosynthesis (ii) uptake and activation, (iii) elongation and desaturation, and (iv) β-oxidation. We found that exhausted CD8^+^ T cells exhibited a distinctive fatty acid transcriptomic profile. Genes associated with *de novo* fatty acid synthesis, including *ACLY*, *ACACA* and *FASN*, were only weakly expressed as compared to CD8^+^ effector and memory T cells, suggesting limited intrinsic lipogenesis (Fig 2E). By contrast, uptake of fatty acid was prominent, with high FABP5 expression (Fig 2F). These cells also showed expression of elongation enzymes, most notably *ELOVL1* and *ELOVL5*, together with late-stage β-oxidation components (*ACAA2*, *ECHS1*, *HSD17B10*, *ETFA*, *ETFB*) (Fig 2, G and H). However, the mitochondrial entry enzyme *CPT1A* and the acyl-CoA dehydrogenase (*ACAD*) family were weakly expressed. This profile suggested that exhausted T cells were capable to activate and elongate exogenous fatty acids, yet they were not efficiently funnelled into complete oxidation or biosynthesis, favouring the accumulation of intermediates and metabolic imbalance.

In contrast, regulatory T cells expressed *ACSL5* and a very high level of *ELOVL5*, suggesting the generation of long-chain polyunsaturated fatty acids that may contribute to membrane remodelling and the production of immunoregulatory lipid mediators (Fig 2G). Although CPT1A expression was low, they displayed elevated *HADHA*/*HADHB* (the mitochondrial trifunctional complex) together with *DECR1* and *ETFA*/*ETFB*. This pattern suggested that regulatory T cells engaged a restrained form of β-oxidation that slowed at the terminal thiolytic step while maintaining downstream oxidative capacity. Unlike exhausted T cells, the accumulation of β-oxidation intermediates was unlikely to cause lipotoxic stress because fatty acid entry into mitochondria was throttled and overall metabolic flux remained low (Fig 2H).

Effector CD8^+^ T cells showed increased expression of *ACLY*, suggesting active *de novo* lipogenesis (Fig 2E). The induction of *CPT1A*, *ACADM* and *ACADVL* indicated that entry into mitochondrial oxidation is at least partially engaged, although downstream enzymes including *HADHA*, *HADHB*, *ACAA2*, *ECHS1*, *HSD17B10* and *DECR1* were weakly expressed, suggesting incomplete oxidation (Fig 2H). Instead, activation pathways were engaged, together with elongation activity, highlighting a bias towards lipid remodelling. Effector memory T cells broadly mirrored this profile but with higher FABP5, indicating greater capacity for exogenous fatty acid uptake (Fig 2F). They also displayed slightly stronger downstream oxidative capacity (*HADHA*), suggesting a shift towards more balanced utilisation of fatty acids to support long-term persistence (Fig 2H). Naive T cells expressed only minimal levels of transport, activation, elongation and oxidation genes. However, they showed a high level of *SREBF1*, suggesting a poised transcriptional state in which lipogenic pathways are primed, consistent with their metabolically quiescent but activation-ready phenotype (Fig 2E).

### Exhausted T cells display partial triglyceride synthesis but limited storage capacity

Triacylglycerol metabolism was categorised into biosynthesis, lipid droplet formation and lipolysis. Exhausted T cells displayed partial engagement of the triglyceride synthesis pathway by expressing high *AGPAT2*, which generates phosphatidic acid (Fig 2I). However, upstream (*GPAT4*) and downstream enzymes (*LPIN*s, *DGAT*s) were largely absent. Expression of droplet-coating proteins such as *PLIN2/3* were also low, suggesting that triglyceride intermediates accumulate without progressing to storage or mobilisation.

Regulatory T cells showed a different pattern, with low expression of triglyceride synthesis genes but high levels of lipase *PNPLA2* together with its co-activator *ABHD5*, as well as *PLIN3*. (Fig 2, J and K). This combination suggests the presence of small, dynamic lipid droplets that can be efficiently hydrolysed to provide fatty acids for oxidation, while also allowing neutral lipids to be stabilised and stored in a controlled manner.

Effector CD8^+^ T cells expressed *GPAT4*, *AGPAT1/2/3* and *LPIN1/2*, indicating production of intermediates up to diacylglycerol (Fig 2, I and J). *DGAT*s were low, suggesting that neutral lipid storage is not a major fate, although a moderate level of *PLIN2* was present, allowing limited droplet stabilisation. Lipolysis was minimal, implying that diacylglycerol is more likely channelled into membrane phospholipid synthesis or retained as a signalling intermediate. Effector memory cells followed a similar profile with strong *LPIN1/2*, but *PLIN2* was low, further reducing their capacity to stabilise neutral lipids for storage. Naive T cells showed weak expression of early genes for triglyceride metabolism, consistent with a metabolically restrained phenotype.

### Exhausted T cells show impaired cholesterol biosynthesis with strong efflux

Cholesterol metabolism was categorised into biosynthesis, uptake and transport, esterification and mobilisation, and efflux. Exhausted CD8⁺ T cells expressed upstream mevalonate pathway genes (*HMGCR*, *PMVK*, *IDI1* and *FDPS*) with moderate expression of downstream sterol enzymes (*SQLE*, *LSS*, *CYP51A1* and others) (Fig 2L). This profile indicates that the sterol biosynthetic machinery is present but expressed at restrained levels. Genes responsible for cholesterol uptake and transport genes (*LDLR*, *STARD4/5* and *NPC1*) as well as esterification (*SOAT1*) were moderately expressed (Fig. 2, M and N). Most strikingly, *ABCG1* was highly expressed, implying active cholesterol efflux and highlighting a state in which synthesis is engaged but accumulation of cholesterol is limited (Fig 2O).

Regulatory T cells displayed robust expression of the upstream biosynthetic branch and particularly high *FDFT1*, committing flux into the sterol pathway (Fig 2L). Expression of *SQLE* and subsequent enzymes was detectable but at low levels, indicating that cholesterol biosynthesis is engaged but not strongly executed. Uptake pathways were largely absent apart from *NPC2*, whereas both *SOAT1* and *LIPA* were expressed, supporting an esterification–hydrolysis cycle that buffers intracellular cholesterol without driving net accumulation (Fig 2, M and N). Efflux transporters were low, indicating that, unlike exhausted CD8⁺ T cells, regulatory T cells retain cholesterol internally and sustain homeostasis through storage and regulated mobilisation rather than continuous synthesis and export (Fig 2O). Effector CD8⁺ T cells expressed moderate levels of upstream biosynthetic genes, similar to exhausted T cells, but additionally showed *SOAT1* together with elevated *LIPA* (Fig 2, L and N). Memory T cells resembled effector cells but with lower *LIPA*, indicating reduced capacity for ester mobilisation. Naive cells expressed minimal biosynthetic genes but a high level *LDLRAP1*, perhaps reflecting a preference for cholesterol acquisition (Fig 2M).

Taken together, T cell subsets in ascites exhibited distinct lipid transcriptomes that may shape their functions. Exhausted CD8⁺ T cells remained metabolically active but accumulated fatty acid and triglyceride intermediates due to incomplete utilisation, while cholesterol synthesis was outweighed by strong efflux, leading to defective lipid homeostasis. On the contrary, regulatory T cells adopted a more controlled lipid transcriptomic programme with restrained fatty acid oxidation and cholesterol retention, maintaining metabolic stability that may support their immunosuppressive function.

### Ascites programmed lipid metabolic state of T cells

Since T cells in ascites might have been primed or activated before they were extruded into the peritoneal cavity, T cells isolated from ascites might not fully represent the impact of ascites on T cells. We next cultured PBMC-derived T cells in acellular ascites fluid to compare their transcriptome with the untreated T cells. Differential expression analysis revealed marked alterations in lipid metabolic pathways. Genes responsible for cholesterol biosynthesis and uptake, including *HMGCS1*, *DHCR24* and *LDLR*, were downregulated, indicating reduced capacity for sterol production and receptor-mediated acquisition (Fig 3. A and B). By contrast, cholesterol efflux transporters *ABCA1* and *ABCG1* were strongly induced, suggesting enhanced export of cellular cholesterol. Fatty acid pathways were also activated, with upregulation of *ACACA* and the lipogenic transcription factor *SREBF1*, together with *CPT1A*, the rate-limiting enzyme for mitochondrial import.

**Fig 3.**
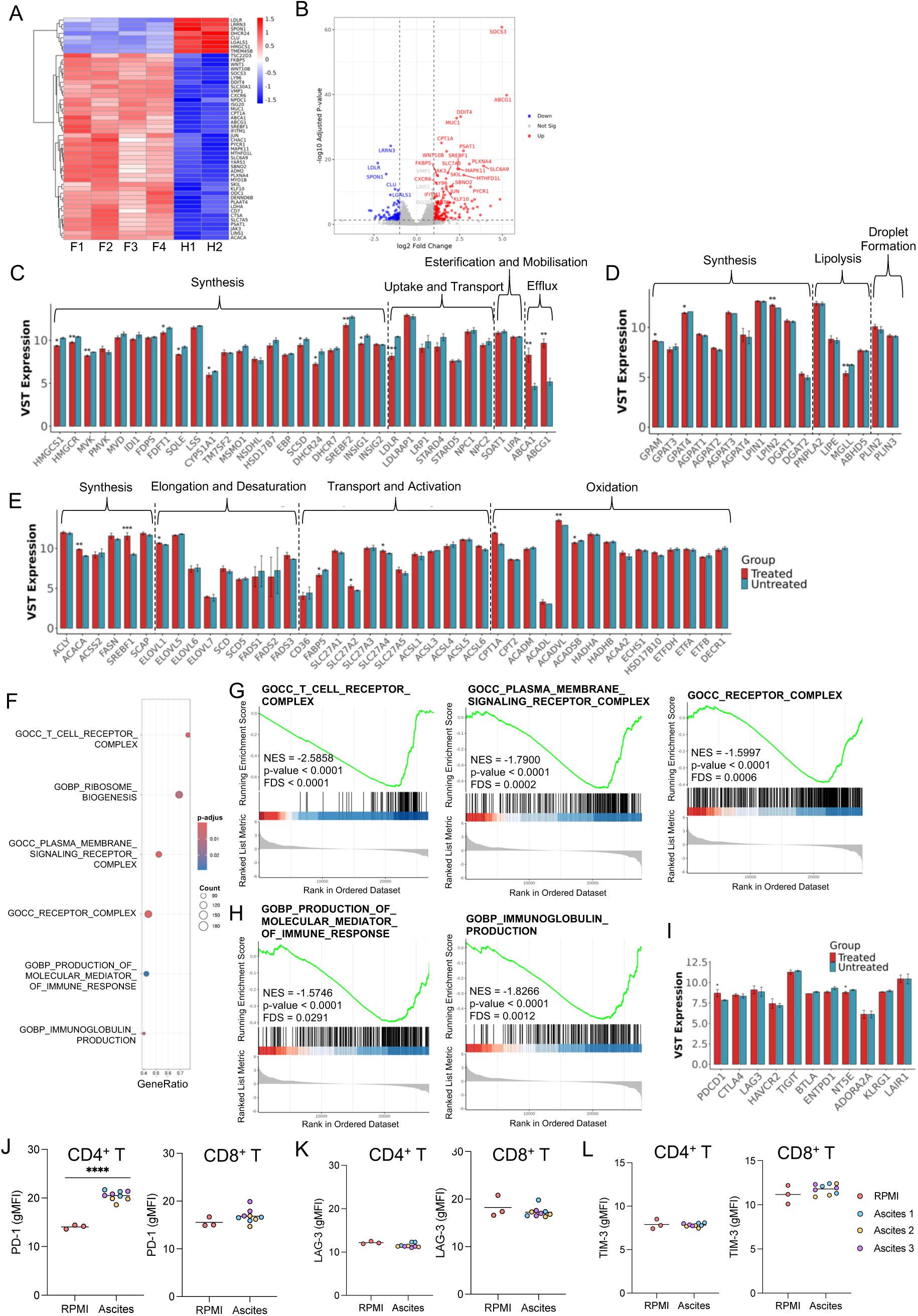
Ascites exposure drives T cells into a metabolically imbalanced, lipid-reprogrammed state with impaired TCR complex signalling. PBMC-derived T cells were co-cultured with ascites (*n* = 4) or serum-supplemented RPMI (*n* = 2) followed by transcriptomics profiling with bulk RNA-sequencing. (A) Heatmap and (B) volcano plot showing top differentially expressed genes in ascites-treated versus untreated T cells. (C to E) Expression of key genes involved in cholesterol metabolism (C), triglyceride metabolism (D), and fatty acid metabolism (E) in ascites-treated and untreated T cells. (F) Dot plot of all significantly enriched gene sets in ascites-treated T cells, as identified by GSEA using MSigDB C5 (Gene Ontology). (G and H) Enrichment plots of representative gene sets related to receptor complex and signalling (G) and production of immune mediators and immunoglobulins (H). (I) Normalised expression of inhibitory receptors in ascites-treated and untreated T cells. (J to L) Flow cytometric analysis of PD-1 (J), LAG-3 (K), and TIM-3 (L) expression on CD4⁺ and CD8⁺ T cells following ascites treatment (*n* = 3). Data are shown as the mean.

The impact of ascites on cholesterol metabolism was further confirmed by a reduction of expression of a range of biosynthetic genes and uptake mediators, together with a strong induction of efflux transporters (Fig 3C). Meanwhile, triglyceride synthesis and lipid formation were mostly unchanged, but *MGLL*, the enzyme responsible for breaking down monoacylglycerol to release free fatty acids, was significantly downregulated (Fig 3D). This implied reduced mobilisation of stored lipids and limited availability of fatty acids for downstream use. Fatty acid metabolism showed a moderate reprogramming, in which *de novo* synthesis was transcriptionally primed (Fig 3E). Importantly, mitochondrial entry genes (*CPT1A*) and early oxidation genes (*ACADVL*) were induced, yet downstream β-oxidation enzymes were not proportionally elevated, perhaps indicating incomplete oxidation.

Together, these data suggested that acellular ascites drove T cells into a metabolically imbalanced state characterised by strong cholesterol efflux and partial engagement of fatty acid oxidation, favouring accumulation of intermediates and metabolic stress.

### Ascites impairs TCR plasma membrane signalling

To understand what biological processes ascites were intervening, we performed GSEA analysis on the “C5: Ontology” signature, and found that ascites has a prominent role in disrupting membrane signalling (Fig 3F). Negative enrichment of “T cell receptor (TCR) complex” ranked top among all, supported by the negative enrichment of “plasma membrane signalling complex” and “receptor complex”, highlighting transcriptomic evidence consistent with impairment of TCR complex signalling (Fig 3G).

GSEA analysis also showed negative enrichment of “production of molecular mediator of immune response” and “immunoglobulin production” in acellular ascites-treated T cells, indicating reduced capacity to generate cytokines, chemokines, and co-stimulatory signals, as well as diminished support for B cell-mediated antibody responses (Fig 3H).

Further gene expression analysis confirmed that immune suppression in ascites-treated T cells is unlikely to be primarily mediated by broad upregulation of inhibitory receptors, as most were unchanged, with *PDCD1* (PD-1) being the only checkpoint showing significant induction upon ascites exposure (Fig 3I). Flow cytometric analysis confirmed acellular ascites fluids upregulated PD-1 expression on CD4^+^ T cells but not the CD8^+^ T cells (Fig 3J). Meanwhile, LAG-3 and TIM-3 expression levels were not affected by ascites treatment (Fig 3, K and L)

### Lipids accumulate in activated T cells in ascites

To test whether T cells would uptake lipid from ascites, we depleted lipids in ascites fluids with an adsorption and clarification reagent, which selectively removes lipids, lipoproteins, floating fats and cell debris from biological samples. Lipid depletion was confirmed by visual clarification of the ascites samples, although a quantitative lipid profiling (e.g., by mass spectrometry or colorimetric lipid assays) was not performed for assessing the depletion efficiency. We cultured PBMC-derived T cells in either serum-supplemented RMPI or ascites or lipid-depleted ascites (LD ascites), and found that there was not a significant difference in lipid accumulation in T cells across treatments (Fig 4A).

**Fig 4.**
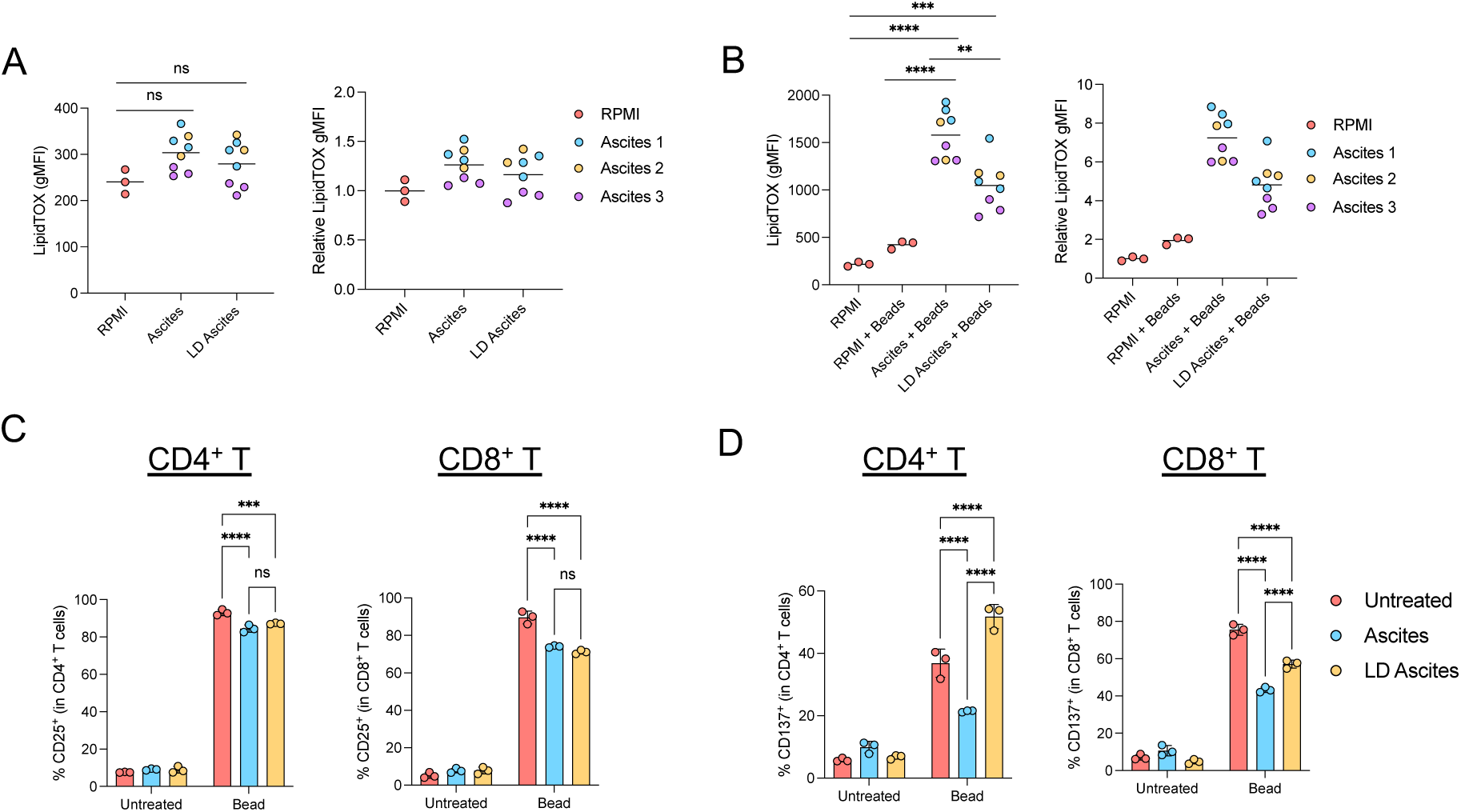
Ascites lipids suppress T cell activation, and lipid depletion selectively rescues CD137 expression. PBMC-derived T cells were cultured in serum-supplemented RPMI, ascites, or lipid-depleted ascites for 72 hours, in the presence or absence of anti-CD3/CD28 Dynabeads. Intracellular lipid levels were assessed by LipidTOX staining. (A) LipidTOX intensity (left) and relative fold change normalised to the untreated control (right), *n* = 3. (B) LipidTOX intensity (left) and relative fold change normalised to unstimulated control (right), *n* = 3. Data are shown as the mean. (C to D) Expression of CD25 (C) and CD137 (D) on CD4⁺ and CD8⁺ T cells, as determined by flow cytometry. Data are shown as mean ± s.d.

We next hypothesised that lipid accumulation would be more pronounced in activated T cells, given their increased demand for metabolic precursors to support proliferation. Following CD3/CD28 co-stimulation, T cells cultured in serum-supplemented RMPI showed a 1.9-fold increase in their lipid content (Fig 4B). Strikingly, T cells cultured in ascites had a 7.2-fold increase in the lipid content, which was partially reduced to a 4.8-fold increase in lipid-depleted ascites. These findings suggested that T cell activation status and the extracellular environment strongly influenced intracellular lipid accumulation.

### Removal of lipid selectively restores CD137 expression on activated T cell

Lipid accumulation has been reported to cause T cell dysfunction (14). We hypothesised that removing the lipid in ascites could at least partially restore the T cell functions. We confirmed that T cells stimulated by CD3/CD28 Dynabeads in ascites were less activated than those in serum-supplemented medium, as demonstrated by the reduction of CD25 expression (Fig 4C). However, lipid depletion in ascites did not restore or increase the CD25 on CD4^+^ and CD8^+^ T cells. Interestingly, lipid depletion selectively restored the level of CD137, which serves as both an activation marker and a co-stimulatory signal (Fig 4D). Meanwhile, non-activated T cells showed no significant changes in CD137 expression across different culture conditions.

The differential effects on CD25 and CD137 likely reflect distinct regulatory mechanisms: CD137 expression depends primarily on the TCR complex, whereas CD25 induction relies more on cytokine-driven pathways such as IL-2 signalling (22,23). Notably, the lipid depletion procedure using Cleanascites may have co-depleted soluble factors such as cytokines, growth factors, or other mediators present in ascites, in addition to lipids. This potential co-depletion of IL-2 and other cytokines could contribute to the failure to restore CD25 expression. However, the selective rescue of CD137, a marker dependent on TCR signalling and lipid raft integrity, aligned with our GSEA analysis that lipid accumulation in ascites disrupted plasma membrane signalling, and is consistent with the hypothesis that lipid removal restores membrane-dependent signalling (24).

### Lipid removal restores bispecific T cell engager-mediated anti-tumour activity

Bispecific T cell engagers (BiTEs) targeting EpCAM (EpCAM × CD3), such as catumaxomab, have been developed in treating malignant ascites (25). Results of clinical trials were encouraging, leading to its initial approval in Europe in 2009 (26–30). Although it was later withdrawn for commercial reasons, catumaxomab has recently regained approval in the European Union for the intraperitoneal treatment of malignant ascites with EpCAM^+^ carcinomas (30). Meanwhile, another EpCAM-targeting BiTE, M701, is currently being evaluated in clinical trials (31,32).

We sought to understand whether depleting lipid could further enhance the anti-tumour effect of the EpCAM BiTE in ascites. By co-culturing PBMC-derived T cells with EpCAM BiTE and ovarian cancer cells, HeyA8, we found that EpCAM BiTE-mediated T cell activation, as measured by both CD25 and CD137, was suppressed in ascites fluid harvested from two different ovarian cancer patients. As expected, lipid removal could not restore the level of CD25 expression (Fig5, A and B). In contrast, CD137 expression on CD4^+^ T cells treated with lipid-depleted ascites was completely restored or even higher (Fig 5C). CD137 expression on CD8^+^ T cells was also partially restored (Fig 5D). Meanwhile, T cells treated with Control BiTE targeting an irrelevant non-human antigen showed no effect on T cell activation across different treatment conditions.

**Fig 5.**
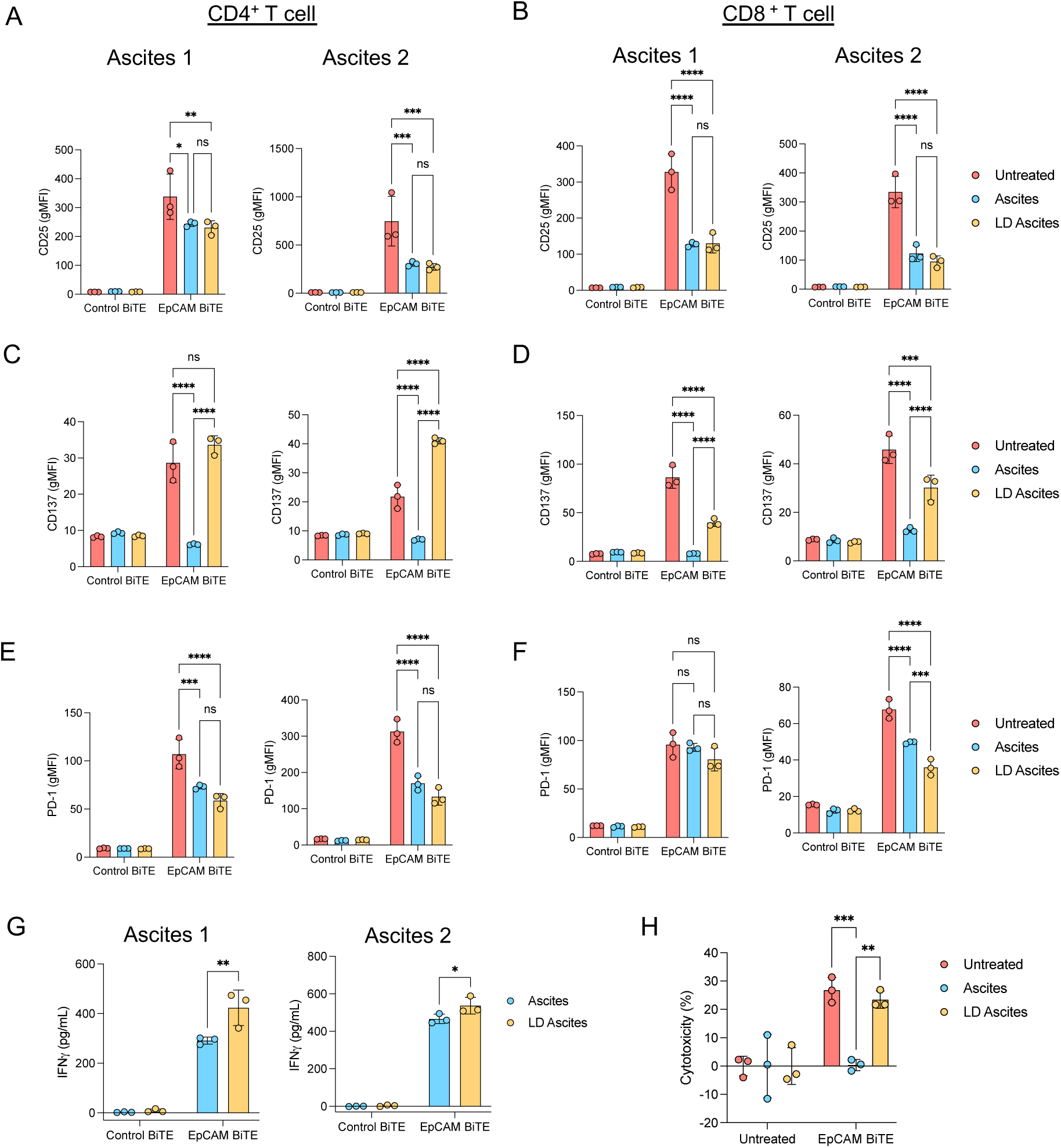
Lipid depletion restores EpCAM BiTE-mediated T cell activation and cytotoxicity in ascites. PBMC-derived T cells were co-cultured with HeyA8 cells and EpCAM BiTE in serum-supplemented RPMI, ascites, or lipid-depleted ascites for 72 hours. Expression of CD25 (A and B), CD137 (C and D), and PD-1 (E and F) on CD4⁺ and CD8⁺ T cells was assessed by flow cytometry. (G) Concentration of IFN-γ in the slice culture medium. (H) T cell-mediated cytotoxicity across treatment groups. Data are shown as mean ± s.d.

The reduction of PD-1 expression on activated T cells in ascites was expected as T cell activation was attenuated. Surprisingly, when we restored T cell activation (CD137) in lipid-depleted ascites, there was a marginal decrease in PD-1 expression, rather than an increase typically associated with enhanced activation (Fig 5, E and F). This unexpected PD-1 behaviour may be explained by increased availability of CD137, whereby engagement with CD137 ligand (CD137L) expressed on cancer cells may ameliorate T cell exhaustion (33). Importantly, from a therapeutic perspective, this dissociation may be beneficial, as T cell activation without concomitant PD-1 upregulation may result in effector cells that are less susceptible to PD-L1-mediated suppression. In addition, T cells activated in lipid-depleted ascites also exhibited a more pro-inflammatory phenotype, with increased secretion of IFN-γ (Fig 5G). The potential of lipid depletion in restoring T cell function was further validated in a cytotoxic killing assay, where ascites abolished EpCAM BiTE-mediated cytotoxicity, which was almost fully rescued by lipid removal (Fig 5H). Collectively, these data suggested that lipid depletion could restore T cell activation, thereby restoring BiTE-mediated cytotoxicity in ascites.

## DISCUSSION

Ascites is a well-recognised immunosuppressive niche present in nearly all cases of ovarian cancer metastases (34). We showed that altered lipid metabolism is one of the key drivers of T cell dysfunction in ovarian cancer ascites. We demonstrated that acellular ascites fluids induced transcriptomic changes in lipid metabolism in T cells, characterised by incomplete β-oxidation and enhanced cholesterol efflux, resembling the transcriptomic profile of exhausted T cells. This likely results in the accumulation of lipid intermediates and a state of metabolic imbalance, making T cells vulnerable to lipotoxic stress and impairing their effector functions.

Importantly, however, ascites-treated T cells did not express high levels of inhibitory receptors. Instead, they exhibited disrupted plasma membrane TCR signalling, which was associated with suppressed T cell activation. In other words, the altered lipid metabolic state may raise the threshold for T cell activation, driving them into a hyporesponsive or anergy-like state rather than classical exhaustion (35).

Cholesterol imbalance is believed to be one of the key factors that dampen the TCR signalling (36). We showed that exhausted T cells and ascites-treated PBMC-derived T cells both demonstrated strong induction of cholesterol efflux transporters along with weak commitment to sterol biosynthesis. This pattern perhaps suggests an activation of the LXR-ABCG1 protective programme to prevent the accumulation of free cholesterol and lipid intermediates, which could otherwise be cytotoxic (37). While such efflux may protect cellular viability, it comes at a cost: depletion of plasma membrane cholesterol and destabilisation of lipid rafts (38). These cholesterol-rich microdomains are essential for TCR clustering and for the recruitment and activation of proximal kinases such as Lck and ZAP-70 (16,38). Disruption of rafts by reduced membrane cholesterol may therefore explain the attenuated TCR complex signalling seen in exhausted or ascites-treated T cells.

*Abcg1*-deficient T cells have been reported to accumulate more plasma membrane cholesterol and increase lipid raft formation to make T cells primed for stimulation (39). In addition, inhibiting cholesterol esterification via ACAT1 blockade has been reported to increase plasma membrane free cholesterol, thereby restoring TCR signalling and reinvigorating CD8⁺ T cell effector function in tumours (16). These models provide a plausible explanation for the selective restoration of CD137 (4-1BB) but not CD25 (IL-2Rα) after lipid depletion. CD137 expression is heavily dependent on strong TCR/CD28 signalling, which requires intact lipid rafts. Restoration of membrane cholesterol may re-enable TCR clustering and the downstream pathway needed to induce CD137 (22). By contrast, CD25 induction depends additionally on sustained cytokine IL2-mediated signalling, which is less likely to be rescued by repairing membrane cholesterol or raft structure (23). Indeed, deletion of *ABCG1* genes in T cells did not show an increase in IL-2 secretion (39). Thus, we proposed that cholesterol efflux in the ascites environment imposed a selective impairment of TCR complex-driven activation, leading cells into a state more akin to anergy or hyporesponsiveness rather than classical exhaustion.

We also showed that exhausted T cells exhibited a different lipid transcriptomic profile than regulatory T cells in ascites, giving an explanation of why ascites do not abrogate the immunosuppressive functions of regulatory T cells (40). We found that regulatory T cells adopted a more coherent lipid programme, characterised by robust lipid droplet turnover and restrained β-oxidation,. This controlled lipid utilisation supported sustained mitochondrial metabolism, providing not only ATP but also acetyl-CoA, which could post-translationally modify FoxP3 through acetylation to protect FoxP3 from degradation (41,42). This may explain how regulatory T cells maintained their immunosuppressive phenotype in the lipid-rich ascites environment.

These mechanistic insights established a compelling rationale for therapeutic intervention in a lipid-depleted environment. Bispecific T cell engagers (BiTEs), in particular EpCAM x CD3, have shown promising efficacy in clinical trials for treating malignant ascites and improving puncture-free survival (28,32). More recently, preclinical BiTEs targeting antigens such as MUC16, and AXL were under evaluation for ovarian cancers and showed potential in ascites models (43,44). Importantly, our findings indicated that metabolic reprogramming via lipid depletion could restore CD137 expression and enhance BiTE-mediated cytotoxicity, suggesting that combining BiTEs with a lipid-targeting approach might significantly enhance anti-tumour T cell activity.

In addition to that, anti-CD36 monoclonal antibody, PLT012, that blocks lipid uptake to reprogram the tumour microenvironment is under development. By inhibiting CD36-mediated fatty acid transport, PLT012 depletes lipid-dependent immunosuppressive cells while activating antitumour CD8^+^ T cells and NK cells, demonstrating synergistic activity with checkpoint inhibitors in preclinical models (45). This therapeutic strategy validates the broader concept that targeting lipid metabolism can enhance T cell-mediated antitumour immunity, supporting the rationale for combining lipid-targeting approaches with BiTEs in metabolically challenging environments such as malignant ascites.

While our findings highlighted lipids as a key driver of T cell dysfunction in ascites, it is important to recognise that ascites is a very heterogeneous environment. Beyond altered nutrient availability, it contains different cytokines, growth factors, stromal elements, and immune populations (46). Cancer-associated fibroblasts, tumour-associated macrophages, and myeloid-derived suppressor cells are all abundant in ascites and secrete immunosuppressive cytokines such as TGF-β, IL-10, and VEGF, which may synergise with metabolic stress to dampen effector function (47). Thus, lipid accumulation represents only one axis of suppression, and additional factors likely act in concert to impose a multilayered barrier to effective T cell immunity.

All ascites samples were collected from patients with advanced-stage (FIGO III–IV) high-grade serous ovarian cancer at the time of primary or interval debulking surgery, but some patients had received neoadjuvant chemotherapy prior to ascites collection. The present study was not powered to evaluate the impact of prior treatment on ascites lipid composition or T cell function. Furthermore, our conclusion on lipid metabolism in T cells are primarily derived from transcriptomic data, which reflect gene expression programmes but do not directly confirm protein-level metabolic changes. Functional validation, including lipidomic profiling of ascites composition and proteomic assessment of key metabolic enzymes, would be essential to confirm whether the observed transcriptomic signatures translate into functional metabolic changes.

Overall, we demonstrated that ascites reprogrammed T cell metabolism, characterised by enhanced cholesterol efflux and impaired fatty acid oxidation. Lipid depletion selectively restored CD137 expression and enhanced anti-tumour activity, highlighting the potential of targeting lipid metabolism as a therapeutic strategy in cancer.

## Data and materials availability

The RNA-seq data generated from this study can be obtained from the NCBI GEO database with accession number GSE310228. All data needed to evaluate the conclusions are present in the paper or supplemental materials. Data are available upon reasonable request.

## Acknowledgments

The authors thank all the donors who generously provided blood samples and the patients who consented to the use of their ascites for this research. Their contributions were invaluable to this study.

## Author contributions

Conceptualisation: PKTW, GA, LWS. Methodology: PKTW, GA, LFO. Investigation: PKTW, GA. Visualization: PKTW. Funding acquisition: KF, LWS. Project administration: PKTW, GA. Supervision: KF, LWS. Writing – original draft: PKTW. Writing – review & editing: PKTW, GA, KF, LWS.

---

**Figure S1.**
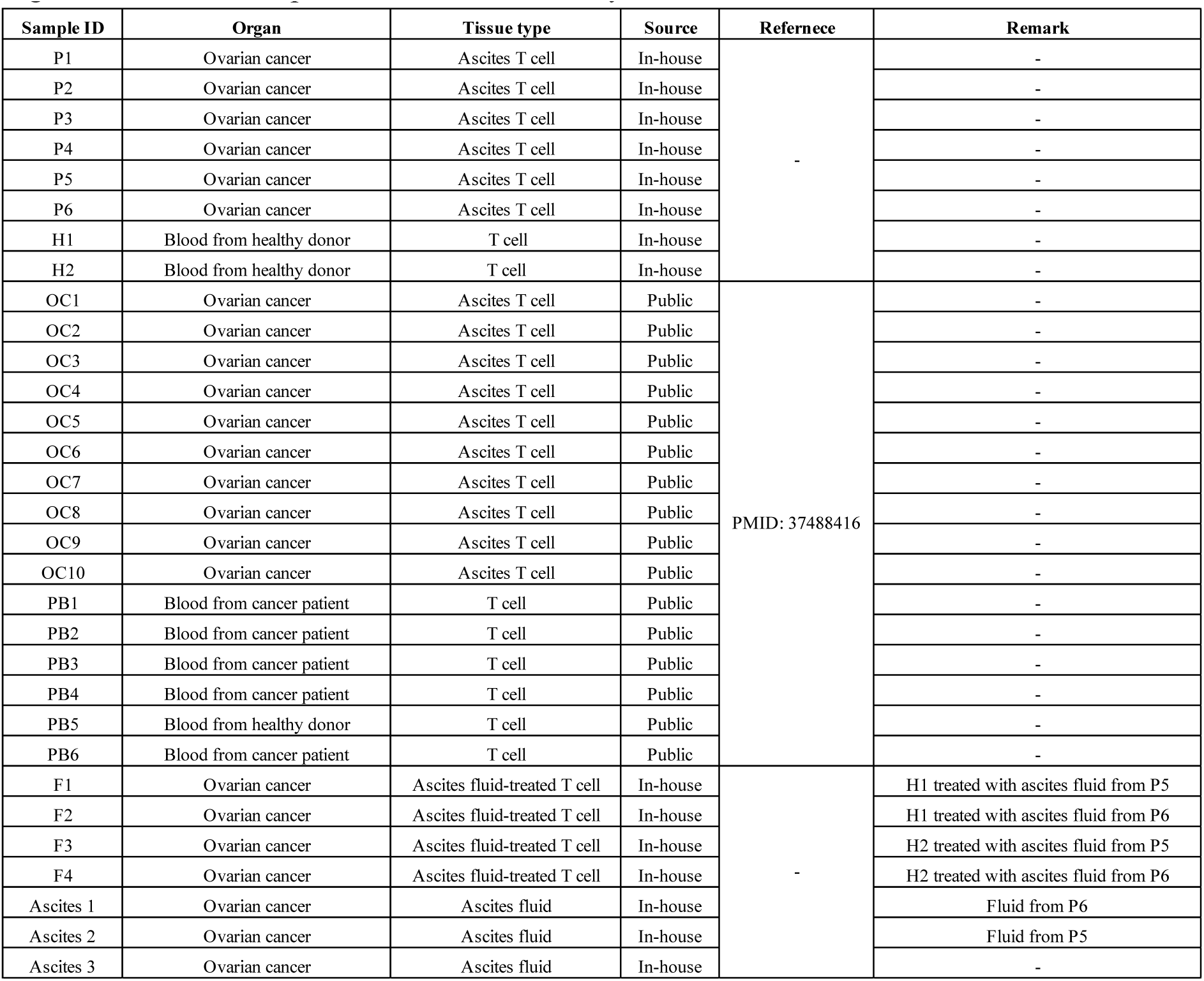
Patient samples included in this study

